# Zbtb24 binding protects promoter activity by antagonizing DNA methylation in mESCs

**DOI:** 10.1101/858662

**Authors:** Haoyu Wu, David San Leon Granado, Maja Vukic, Kelly K.D. Vonk, Cor Breukel, Jihed Chouaref, Jeroen F.J. Laros, Lucia Daxinger

## Abstract

DNA methylation is a key epigenetic modification essential for normal development. How particular factors control DNA methylation patterns and activity of a given locus is incompletely understood. The zinc finger protein Zbtb24 has been implicated in transcriptional activation/repression and the DNA methylation maintenance pathway. Here, using whole genome bisulfite sequencing in mouse embryonic stem cells, we report that besides a general trend towards DNA hypomethylation, many genomic sites gain methylation in the absence of Zbtb24 and they include promoters of actively transcribed genes. DNA hypomethylation is not generally associated with gene expression changes, suggesting that additional epigenetic safeguards are in place that ensure silencing of the affected loci. Remarkably, we identify a set of genes that is particularly susceptible to Zbtb24 occupancy. At these sites, Zbtb24 binding is not only required for gene activity but also required for maintaining the unmethylated state of the promoter.

## INTRODUCTION

DNA methylation is a key epigenetic modification essential for normal development (Smith and Meissner 2013). DNA methylation has been associated with long-term transcriptional repression and in particular methylation of repetitive sequences correlates with gene silencing (Reik 2007). Gene body methylation, which has been linked to active transcription has been suggested to affect gene expression patterns for example, through influencing transcriptional elongation (Baubec et al. 2015), alternative splicing (Lister et al. 2009), or through preventing aberrant transcription initiation (Neri et al. 2017). CpG islands (CGI) are usually free of DNA methylation (Illingworth and Bird 2009). Throughout development, DNA methylation is important to preserve cell type identity and to regulate tissue-specific gene expression patterns (Law and Jacobsen 2010; Greenberg and Bourc’his 2019). However, it remains largely unknown how particular factors control DNA methylation patterns and activity of a given locus, which is in part due to the fact that members of the epigenetic machinery rarely recognize specific DNA sequences. Cross-talk between DNA methylation and transcription factors (TFs) has been suggested. TFs can bind specific DNA motifs, and TF binding can influence the un-methylated state of CpG islands (Brandeis et al. 1994; Thomson et al. 2010; Lienert et al. 2011; Boulard et al. 2015). Conversely, DNA methylation can influence TF binding to their recognition motif (Mann et al. 2013; Domcke et al. 2015; Strogantsev et al. 2015; Yin et al. 2017).

Zinc finger and BTB domain containing 24, ZBTB24, is an enigmatic member of the ZBTB family of C2H2 transcription factors (Lee and Maeda 2012). ZBTB24 contains a BTB domain, an AT-hook motif and eight C2H2-zinc finger domains (Edgar et al. 2005), and is expressed ubiquitously, albeit at relatively low levels (Thompson et al. 2018). In the mouse, Zbtb24 is critical for normal embryonic development and *Zbtb24*^*ΔBTB/ΔBTB*^ mutant mice display prenatal lethality (Wu et al. 2016). Recessive mutations in *ZBTB24* cause Immunodeficiency, Centromeric instability, Facial anomalies (ICF) syndrome (de Greef et al. 2011; Cerbone et al. 2012; Nitta et al. 2013), a disorder characterized by hypomethylation of repetitive DNA (Velasco and Francastel 2019). Aberrant DNA methylation has been reported in ICF patients carrying *ZBTB24* nonsense mutations (Velasco et al. 2018), upon shRNA-mediated ZBTB24 depletion in HCT116 cells, a human colon cancer cell line (Thompson et al. 2018), and in a zebrafish *zbtb24* knock out model (Rajshekar et al. 2018). A combined function of ZBTB24 and DNMT3B has been suggested for mediating some gene body methylation in HCT116 cells (Thompson et al. 2018), but the mechanisms by which ZBTB24 influences DNA methylation at other genomic regions remain unclear. ZBTB24 also functions as a transcriptional activator/repressor, and with its C2H2 zinc finger domain, ZBTB24 can bind to DNA in a sequence-specific manner (Thompson et al. 2018; Aktar et al. 2019; Ren et al. 2019). Interestingly, in mouse, human, and zebrafish ZBTB24 controls expression of cell division cycle associated 7, (CDCA7) (Wu et al. 2016; Rajshekar et al. 2018; Thompson et al. 2018; Velasco et al. 2018; Qin et al. 2019; Unoki et al. 2019), which is another gene that can be mutated in ICF syndrome (Thijssen et al. 2015).

Here, we aimed to establish causal relationships between genome-wide DNA methylation, Zbtb24 binding and gene activity. Using mouse embryonic stem cells (mESCs) knock out for *Zbtb24*, we report integrative genome-wide analyses of DNA methylation, histone modification, TF binding and gene expression profiles. Besides hypomethylation of CpG poor, gene poor regions and gene clusters, we find many sites that gain DNA methylation in the absence of Zbtb24 and they include promoters of actively transcribed genes. Interestingly, we identify a selected group of genes that are particularly susceptible to Zbtb24 occupancy. At those sites, Zbtb24 binding is not only required for promoter activity but also involved in maintaining the unmethylated state of the CGI. In contrast, DNA hypomethylation is not generally associated with gene expression changes, suggesting that additional epigenetic safeguards are in place that ensure silencing of the affected loci.

## RESULTS

### WGBS and RRBS identify hypo- and hyper-DMRs in *Zbtb24* knock out mESCs

To investigate the role of Zbtb24 in DNA methylation pathways we used serum cultured mESC lines derived from *Zbtb24*^*+/+*^ and *Zbtb24*^*ΔBTB/ΔBTB*^ mice (Wu et al. 2016), or *Zbtb24* knock out lines (*Zbtb24*^*-/-*^) generated through CRISPR/Cas9 genome editing in E14 mESCs (**Figure S1A-B**). Whole genome bisulfite sequencing (WGBS) revealed an average global DNA methylation level of ∼75% in wild type (WT) cells, whereas methylation levels significantly dropped (P<1×10^−314^) to around 70% in *Zbtb24*^*-/-*^ cells (**Figure 1A-B**). Particularly distal intergenic regions lost methylation in *Zbtb24*^*-/-*^ cells. Furthermore, introns and regions >2kb downstream of the transcription end site (TES) were associated with reduced methylation levels (**Figure S1D**). Interestingly, we also noticed DNA hypermethylation in *Zbtb24*^*-/-*^ mESCs. The percentage of CpGs with a methylation level >95% was higher in *Zbtb24*^*-/-*^ (18%) when compared to WT (11%) (**Figure S1C**). Similarly, a modest but significant global hypermethylation (P<2.2×10^−16^, one sided Wilcoxon rank sum test for paired samples) was found in reduced representation bisulfite sequencing (RRBS) datasets generated from the independent *Zbtb24*^*+/+*^ and *Zbtb24*^*ΔBTB/ΔBTB*^ mESC lines (**Figure 1D-E; Figure S1B**).

**Figure 1.**
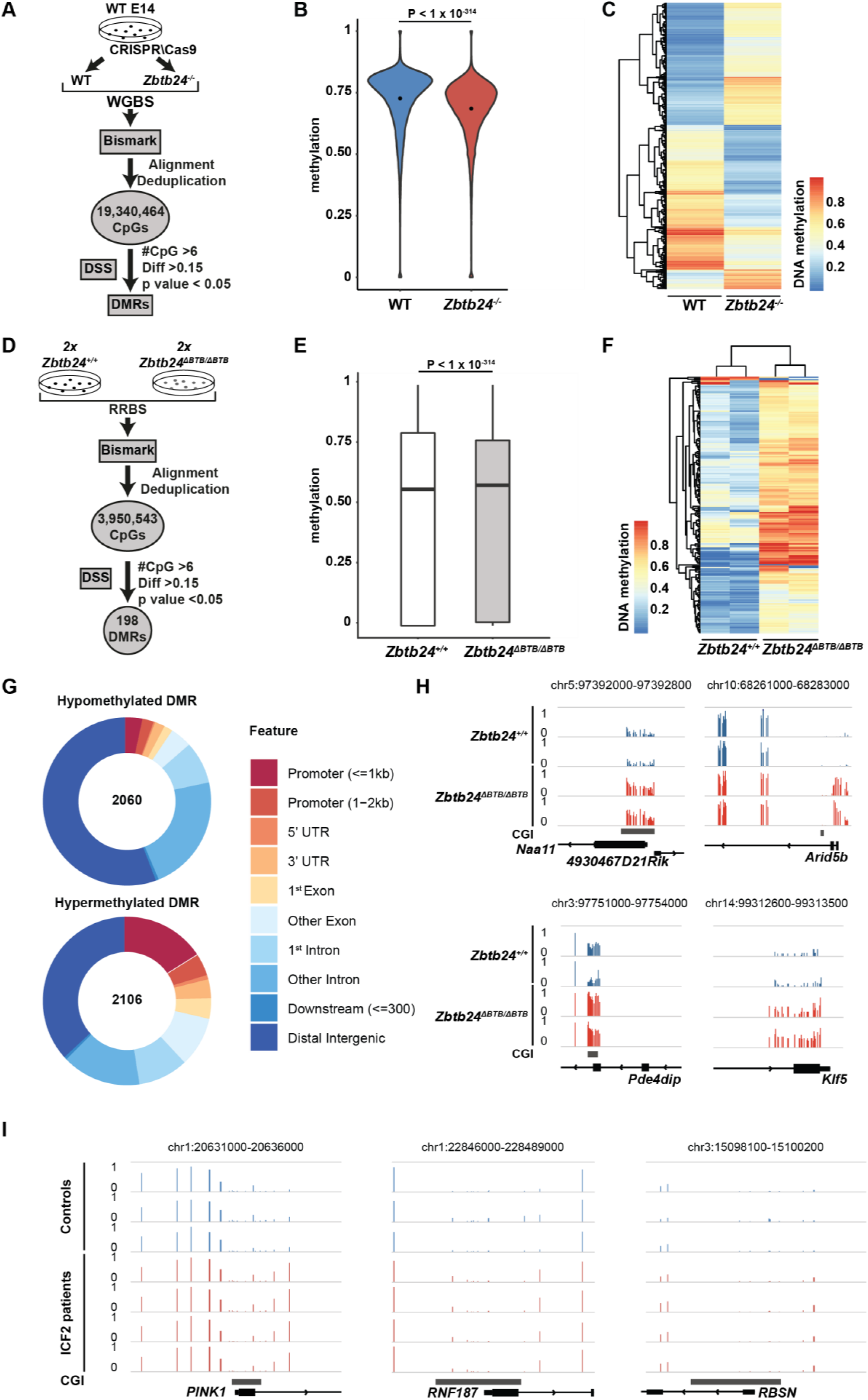
DNA methylation changes in *Zbtb24* mutant mESCs. (A) Schematic depicting the generation of the *Zbtb24*^*-/-*^ mESC line and the WGBS analysis workflow (n = 1 WGBS dataset for each genotype). (B) Violin plot showing global methylation levels (3kb tiles) in WT (blue) and *Zbtb24*^*-/-*^ (red) E14 WGBS samples. The *y*-axis represents the ratio between methylated Cs and total coverage. Black dots represent the mean. The p-value (P) was calculated with a one side paired Wilcoxon rank sum test. (C) Unsupervised hierarchical clustering and heatmap showing average methylation levels of differentially methylated regions (DMRs) in WT and *Zbtb24*^*-/-*^ mESCs. (D) Schematic showing the RRBS analysis workflow. (n = 2 RRBS datasets for each genotype). (E) Box plot showing global methylation levels in *Zbtb24*^*+/+*^ and *Zbtb24*^*ΔBTB/ΔBTB*^ RRBS samples. DNA methylation levels per CpG were calculated as the average of the two replicates. The p-value (P) was calculated with a one side paired Wilcoxon rank sum test. (F) Unsupervised hierarchical clustering and heatmap indicating average methylation levels of *Zbtb24*^*+/+*^ and *Zbtb24*^*ΔBTB/ΔBTB*^ DMRs. (G) *Donut plot* depicting the distribution of Zbtb24-hypo and -hyperDMRs of different genomic features. The number in the donut hole represents the number of identified DMRs ((≥15% methylation change, coverage of >6 CpGs). (H) UCSC screenshots showing RRBS data for four representative loci with methylation gain in *Zbtb24*^*ΔBTB/ΔBTB*^ mutant mESCs. The *y*-axis indicates the DNA methylation level. (I) Examples of CGI promoters that show differentially methylated CpGs between controls (blue) and ICF2 patients (red). Reanalysed Illumina 450K data from (Velasco et al. 2018).

When compared to wild type, we found 4168 statistically significant differentially methylated regions (DMRs) (p-value<0.05) in *Zbtb24*^*-/-*^ cells using a threshold of ≥15% methylation change and a coverage of >6 CpGs for the WGBS dataset. About half of the Zbtb24-DMRs (n=2060) were hypomethylated (Zbtb24-hypoDMRs), and the other half (n=2106) were hypermethylated (Zbtb24-hyperDMRs) (**Figure 1C; Table S1**). In the RRBS data, which is generally enriched for CpG rich regions (Gu et al. 2011), we found a total of 198 DMRs (≥15% methylation change, coverage of >6 CpGs). The vast majority, n= 195, showed hypermethylation in *Zbtb24*^*ΔBTB/ΔBTB*^ mutants (**Figure 1F; Table S1**).

Consistent with a function for Zbtb24 in maintaining DNA methylation at intergenic regions and gene clusters (Velasco et al. 2018), hypoDMRs encompassed members of large gene families including the clustered protocadherin, olfactory receptor and vomeronasal receptor genes and many hypoDMRs mapped to distal intergenic regions (**Figure 1G; Figure S1E; Table S1**). This also suggests that these targets are conserved between mouse and human. DNA methylation levels of interspersed repeat elements such as Satellites, Lines, ERVs, and SINEs were only modestly reduced in *Zbtb24*^*-/-*^ cells (**Figure S1F**).

In both the WGBS and RRBS datasets, we found many Zbtb24-hyperDMRs in transcription start sites (±2kb) and introns and exons (n=1333 (P= 1.8×10^−96^) WGBS (**Figure 1G; Table S1**) and n=159 RRBS Zbtb24-hyperDMRs (**Table S1**)). Some examples are shown in (**Figure 1H**). Intriguingly, promoters with aberrant DNA methylation gain included Zbtb24 targets previously identified in mESCs, such as *Cdca7* and *Arid5b* (Wu et al. 2016; Ren et al. 2019) (**Table S1**). We also confirmed ZBTB24-associated promoter hypermethylation by analysis of published Illumina 450K datasets from controls and ICF2 patients (Velasco et al. 2018) (**Figure 1I; Figure S2**), and again, some of these sites are known ZBTB24 targets in HCT116 cells (Thompson et al. 2018). We conclude that the effect of Zbtb24 depletion on the mESC methylome is widespread and not limited to DNA hypomethylation. Rather, the promoter methylation gain at known Zbtb24 targets suggests a putative direct link between Zbtb24 occupancy and aberrant DNA methylation state.

### Zbtb24 occupancy and sites of DNA methylation gain are coupled

Local binding of transcription factors can contribute to maintain hypomethylated states of CGI (Krebs et al. 2014). Since Zbtb24 is often found at unmethylated CpG rich promoters (Thompson et al. 2018; Ren et al. 2019), we considered that promoter methylation gain could be a consequence of loss of Zbtb24 binding. In the absence of a ChIP-seq suitable endogenous Zbtb24 antibody we used Ty1-tagged Zbtb24 to investigate genome-wide binding in mESCs (**Figure 2A**). Consistent with previous studies (Thompson et al. 2018; Ren et al. 2019), we found that most Zbtb24 peaks localized to promoters, exons and introns, and some overlapped with intergenic regions (**Figure 2A; Table S2**). The promoters of *Cdca7, Ostc, Rnf187* and *Cdc40* were among the strongest peaks (**Table S2**). Motif enrichment analysis using MEME-ChIP showed that a Zbtb24 motif very similar to a motif recently identified in human HCT116 cells and mESCs (Thompson et al. 2018; Ren et al. 2019), was enriched in our dataset (**Figure 2B; Table S2**).

**Figure 2.**
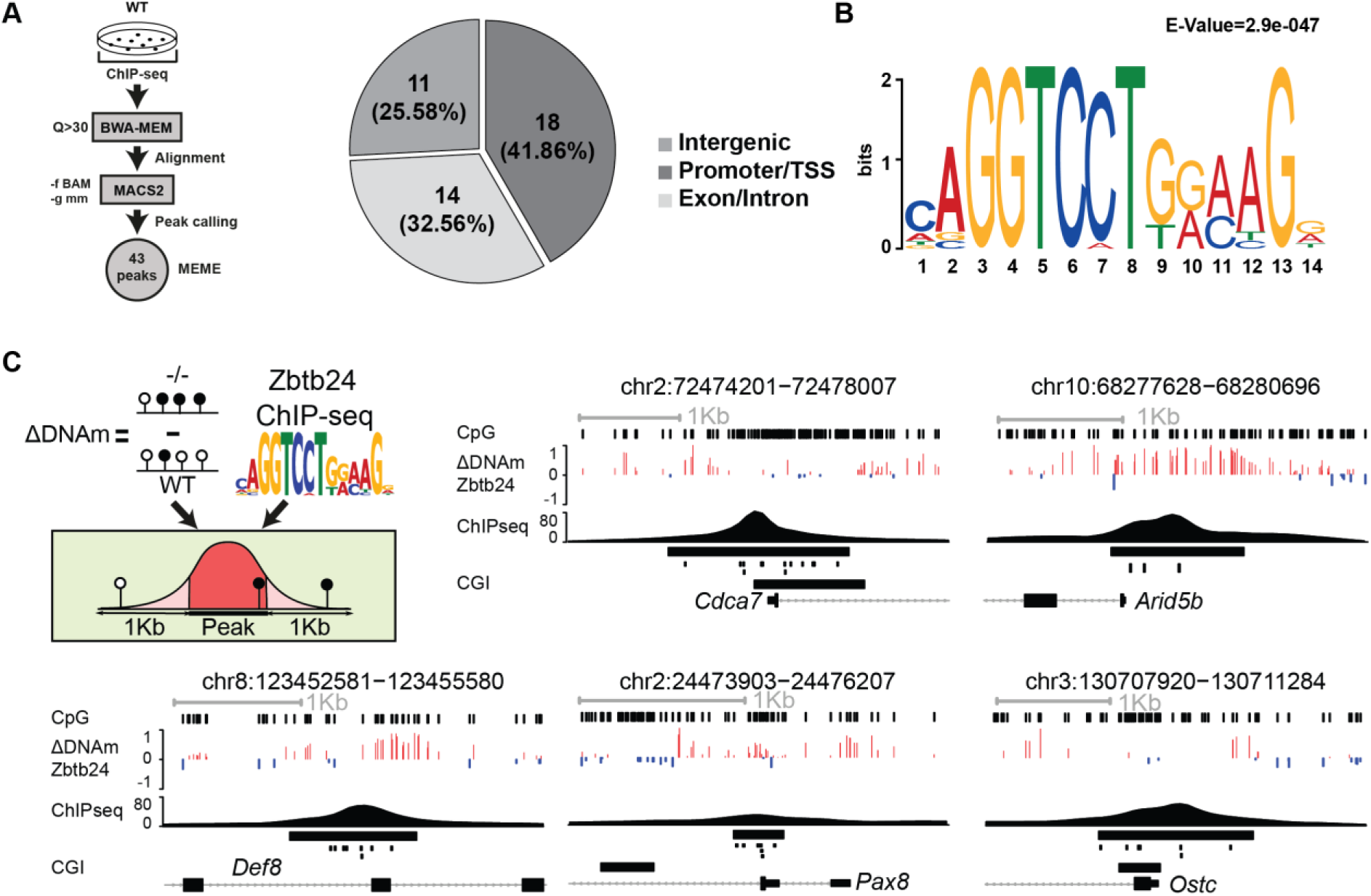
Zbtb24 binding sites overlap with Zbtb24-hyperDMRs. (A) Schematic of the Ty1-Zbtb24 E14 mESC ChIP-seq experiment and analyses pipeline, and genome-wide distribution of identified Ty1-Zbtb24 peaks. The promoter/TSS annotation corresponds to TSS ±2kb. (B) Zbtb24 binding motif in mESCs as identified by MEME (*e*-value in the top right corner). (C) Schematic of DNA methylation, ChIP-seq and histone modification dataset integration, and genome browser screenshots of representative sites where Zbtb24 binding sites (±1kb) overlap with Zbtb24-hyperDMRs. Differential methylation levels (ΔDNAm Zbtb24) between *Zbtb24*^*-/-*^ and WT are depicted in red (positive values) and blue (negative values). The larger black rectangles under the Zbtb24 ChIP-seq peak represent the called peak, small black rectangles represent Zbtb24 motifs.

To assess whether sites of methylation gain in *Zbtb24*^*-/-*^ cells coincide with Zbtb24 peaks, we computed differences in DNA methylation levels between wild type and *Zbtb24*^*-/-*^ cells at Zbtb24 bound sites (±1kb). Indeed, we found 29 Zbtb24-bound sites that acquired DNA methylation in *Zbtb24*^*-/-*^ cells (multiple Zbtb24 peaks within a 1kb region were considered as one peak). These sites included conserved Zbtb24 targets such as the promoters of *Cdca7, Arid5b* and *Ostc* (**Figure 2C; Table S2**). On average, DNA methylation levels around Zbtb24 peaks (±1kb) increased by 7% upon loss of Zbtb24 binding, with some regions gaining up to 34% methylation on average) (**Table S2**). We thus concluded that although a substantial proportion of Zbtb24-hyperDMRs may be attributable to indirect effects, there is a set of genomic sites where promoter methylation seems to occur in response to loss of Zbtb24 binding.

### Gene expression changes upon loss of Zbtb24

The local chromatin environment can influence transcriptional activity and may influence downstream consequences of Zbtb24-related aberrant DNA methylation. Therefore, we next compared Zbtb24-DMRs with published datasets for various histone modifications (Marks et al. 2012). Heat map clustering revealed that Zbtb24-hypoDMRs overlapped with H3K4me3, H3K9me3 and H3K26me3. Zbtb24-hyperDMRs showed strong H3K4me3 enrichment (n=1074 hyperDMRs overlapped with this modification) and some enrichment for H3K36me3 and H3K27me3 (**Figure 3**). Together, this suggests that Zbtb24-related aberrant methylation may affect transcriptionally active loci.

**Figure 3.**
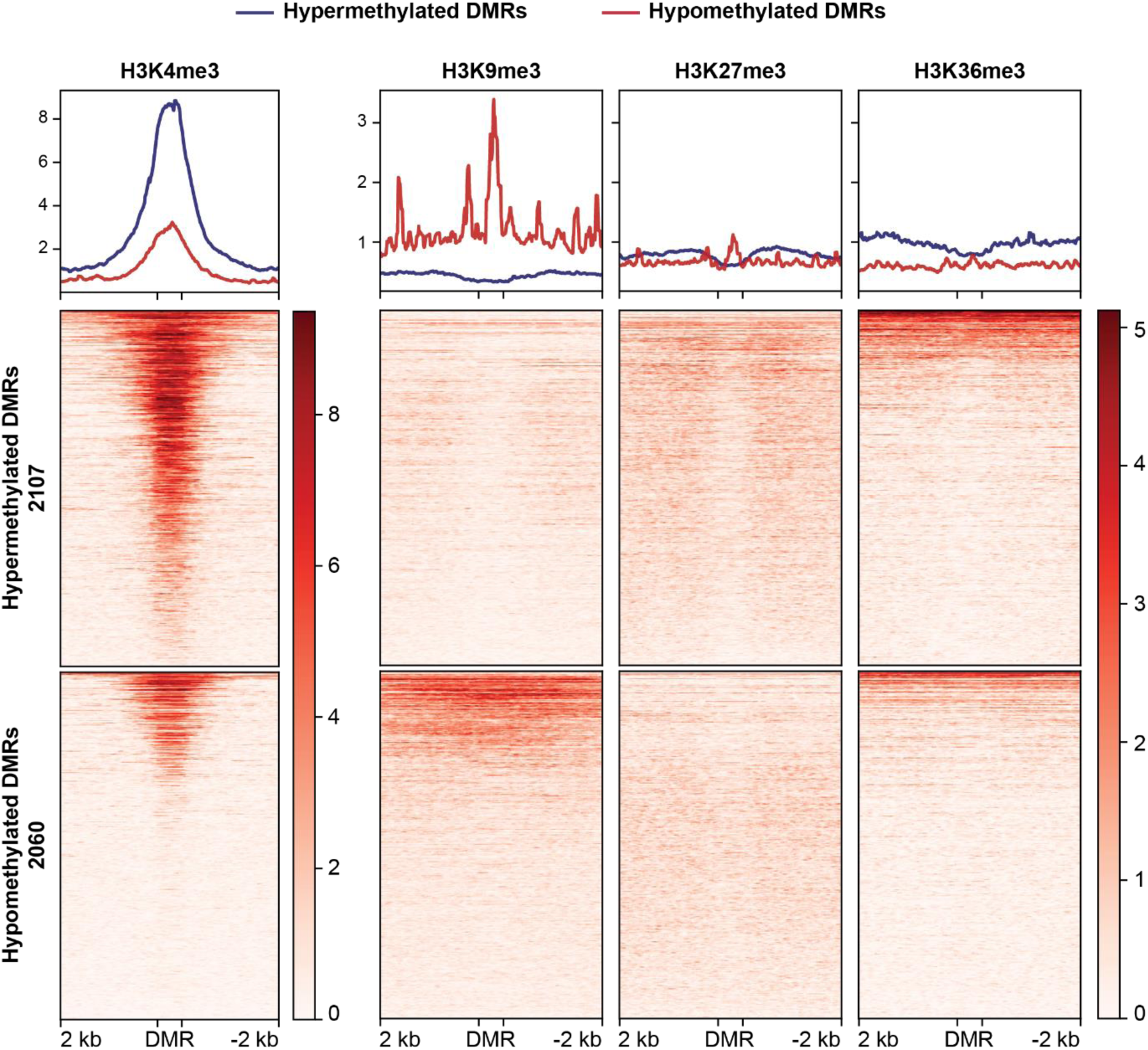
Chromatin environment around Zbtb24DMRs. Histone modification ChIP-seq heatmaps and average profiles centered on Zbtb24-hypo and - hyperDMRs (±2kb). Color intensity represents normalized tag counts.

To examine the functional relevance of Zbtb24-related anomalous DNA methylation we performed total RNA-sequencing (RNA-seq) on wild-type and *Zbtb24*^*ΔBTB/ΔBTB*^ homozygous mutant mESCs grown in serum conditions. In total, 479 genes were significantly differentially expressed between *Zbtb24*^*ΔBTB/ΔBTB*^ and *Zbtb24*^*+/+*^ mESCs (*p*(adj)< 0.05, log_2_FC≥1). 328 genes were upregulated and 151 genes were downregulated in the mutants when compared to wild type controls (**Figure 4A; Table S3**). As expected, well established Zbtb24 targets such as *Cdca7, Rnf187* or *Cdc40* were among the downregulated genes (**Table S3**), consistent with a transcriptional activator function for Zbtb24 at these sites. Of note, although we found that the absence of Zbtb24 was associated with hypomethylation of e.g. the clustered protocadherin genes (**Figure S1E)**, this did not generally correlate with gene expression changes (**Table S3**). We then integrated RNA-seq, Zbtb24 ChIP-seq and WGBS datasets to identify causal relationships between Zbtb24 occupancy, DNA methylation and gene expression levels. Of the 33 Zbtb24 target sites where peaks overlapped with promoters and/or gene bodies, 19 were down-regulated and showed hypermethylation in the regions surrounding the Zbtb24 peak (±1kb) (**Figure 4B**). This suggests that at these sites, loss of Zbtb24 can indeed be causally linked to gene expression change and promoter DNA methylation gain. Of note, in the case of the *Def8* gene, loss of Zbtb24 binding resulted in *de novo* methylation and reduced transcription from an intragenic alternative TSS (**Figure 4C**), providing an example where TF binding influences transcript isoform usage. We also looked for genomic sites where gene expression levels might be affected by promoter DNA hypomethylation (at least 15% methylation change, ≥7CpGs), but found only one such locus (**Figure S3**). Indeed, a number of Zbtb24-hypoDMRs are targets for H3K9me3 (**Figure 3**), which is associated with gene repression and a major silencing pathway in mESCs (Ninova et al. 2019). Therefore, at these Zbtb24-hypoDMRs, persistent H3K9me3 could explain the absence of gene expression changes.

**Figure 4.**
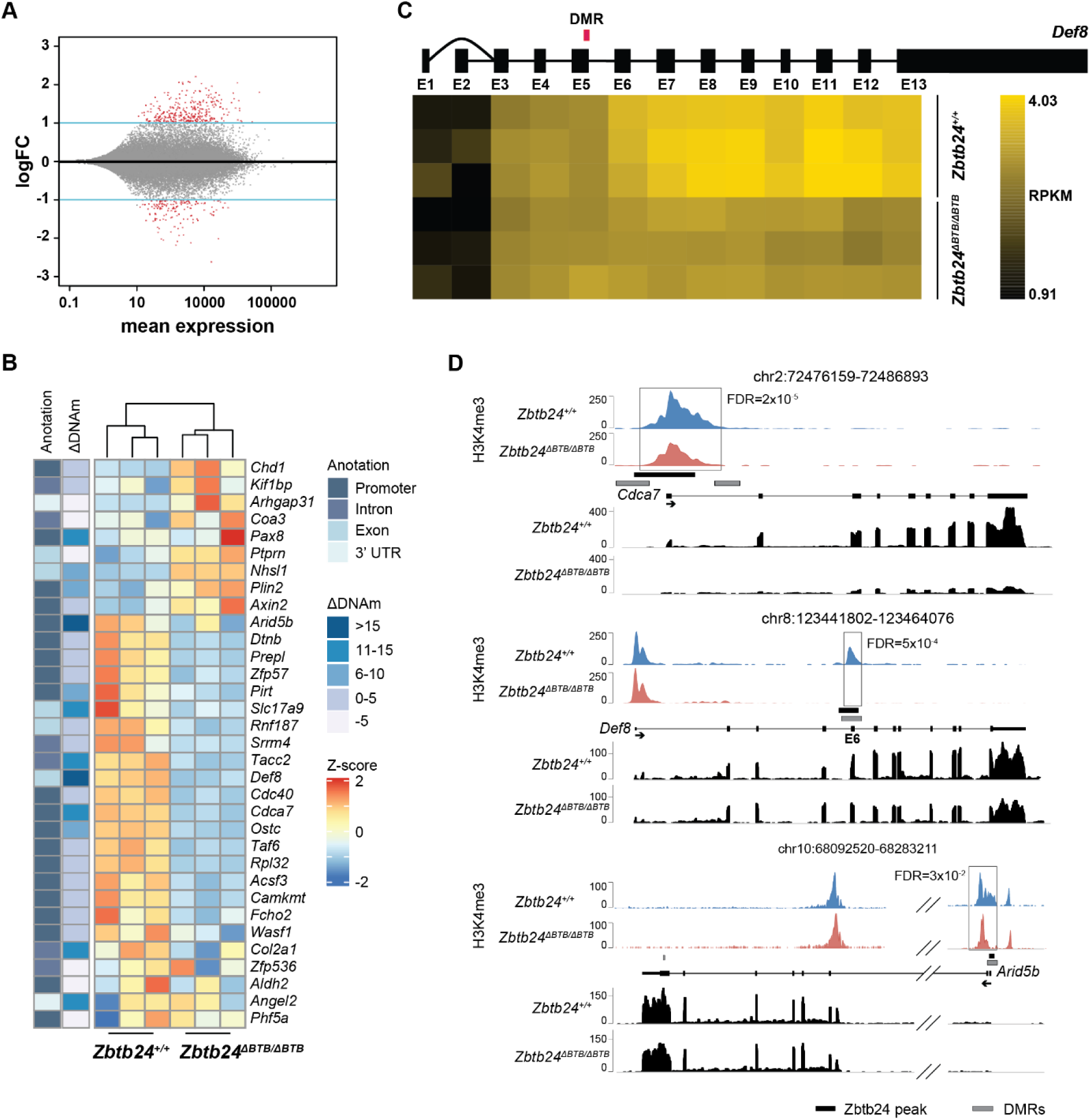
Gene expression levels, DNA methylation and H3K4me3 levels at Zbtb24 bound sites. (A) MA plot of normalized RNA-seq data (n = 3 samples for each genotype). Red dots indicate genes that are differentially expressed between *Zbtb24*^*ΔBTB/ΔBTB*^ and *Zbtb24*^*+/+*^ mESCs (p-adj<0.05; log2FC >1). The *y*-axis shows logFC and the *x*-axis the average log intensity (mean expression). (B) Unsupervised hierarchical clustering and heatmap showing expression levels of Zbtb24 bound sites in *Zbtb24*^*+/+*^ and *Zbtb24*^*ΔBTB/ΔBTB*^ mESCs. Blue denotes lower and red indicates higher expression levels. Gene expression values were normalized by Z-score for each row. The annotation column shows gene features. The ΔDNAm column shows the average differential methylation between WT and *Zbtb24*^*-/-*^ mESCs. (C) Heatmap showing expression values (RPKMs) for the Def8 exons. In *Zbtb24*^*ΔBTB/ΔBTB*^ mESCs, from exon 6 expression is significantly decreased (FDR=6.4×10^−5^), which correlates with a hypermethylated DMR (red rectangle). Dark yellow represents low and light yellow high expression. (D) Representative regions showing H3K4me3 levels, Zbtb24 binding, Zbtb24-hyperDMR position and gene expression levels in *Zbtb24*^*ΔBTB/ΔBTB*^ (red) versus *Zbtb24*^*+/+*^ (blue) mESCs. The H3K4me3 track shows the spike-in normalized ChIP-seq signal (average of two biological replicates). RNA-seq track showing expression levels of proximal genes in *Zbtb24*^*+/+*^ and *Zbtb24*^*ΔBTB/ΔBTB*^ mESCs. Zbtb24^-^ hyperDMRs are represented as grey and Zbtb24 binding sites as black rectangles.

### Residual H3K4me3 helps to preserve CpG island hypomethylation

Visual inspection of Zbtb24 binding sites indicated that DNA methylation gain in *Zbtb24*^*-/-*^ mESCs was restricted to the borders of Zbtb24 peaks, when binding sites overlapped with CGI (**Figure 2C**). Indeed, H3K4me3 can repel DNA (Mikkelsen et al. 2007; Weber et al. 2007; Meissner et al. 2008; Edwards et al. 2010; Noh et al. 2015). It has also been shown that H3K4me3, although considered a signature mark of active promoters, often persists when the gene is inactivated (Guenther et al. 2007; Mikkelsen et al. 2007). Therefore, we reasoned that residual levels of H3K4me3 could prevent spreading of DNA methylation throughout the CGI. To address this, we performed quantitative H3K4me3 ChIP-seq in wild type and *Zbtb24*^*ΔBTB/ΔBTB*^ mutant mESCs (**Figure S3**). Indeed, we found that despite transcriptional downregulation of Zbtb24 target genes, at many promoters and TSSs including *Cdca7* H3K4me3 levels did not change at all or were only modestly decreased in *Zbtb24*^*ΔBTB/ΔBTB*^ mutant mESCs (**Table S4**).

At a handful of places, we observed a strong correlation between decreased H3K4me3 levels and loss of Zbtb24 binding. For example, in the case of *Def8*, Zbtb24 binds in close proximity to an alternative TSS in exon 5 in wild type mESCs. In *Zbtb24*^*ΔBTB/ΔBTB*^ mutants H3K4me3 levels at this binding site were abolished, which was accompanied by a strong DNA methylation gain (**Figure 4D**). Here, Zbtb24 binding appears to be a major determinant for the activity of the locus. Another noteworthy example is the promoter of the long isoform of the *Arid5b* gene, which bears the hallmarks of a poised promoter in that it is marked by both H3K4me3 and H3K27me3 in wild type mESCs. Loss of Zbtb24 binding coincided with DNA methylation gain and H3K4me3 loss (**Figure 4D**). Altogether, these findings suggest that Zbtb24 binding is not only necessary to protect promoter activity but its occupancy can also contribute to maintaining the unmethylated state of CGIs. In its absence, promoters/alternative TSS can become vulnerable to DNA hypermethylation. However, residual H3K4me3 can serve as a safeguard that prevents spreading of DNA methylation across the entire CGI thereby protecting its unmethylated state.

## DISCUSSION

Zbtb24 is a conserved C2H2 zinc finger protein that has been implicated in transcriptional activation/repression and maintenance of DNA methylation. In this study, we have identified a role for Zbtb24 in protecting promoter activity by antagonizing DNA methylation in mESCs. Our WGBS identified regions of DNA hypomethylation in *Zbtb24*^*-/-*^ mutant mESCs, confirming its previously proposed role in a pathway that maintains DNA methylation at gene clusters and intergenic regions (Velasco et al. 2018). However, we did not find clear evidence that DNA hypomethylation is a direct consequence of loss of Zbtb24 binding in mESCs.

Interestingly, a very recent study showed that both the AT-hook and the ZF domain of ZBTB24 are required for heterochromatin localization of the protein but how ZBTB24 gets recruited to these sites remains unclear (Aktar et al. 2019). It has also been shown that Zbtb24 associates with pericentromeric heterochromatin in NIH3T3 cells independent of DNA methylation (Nitta et al. 2013). One possibility is that ZBTB24 is part of a silencing complex that relies on repressive histone modifications and disruption of such a complex could affect DNA methylation patterns. Yet, alternative mechanisms cannot be excluded. Another interpretation of our results could be that Zbtb24-related DNA hypomethylation is a consequence of its interaction with the CXXC-zinc finger protein Cdca7. It has been reported that *Xenopus* Cdca7e interacts with and stimulates the chromatin remodeling activity of Hells (Jenness et al. 2018), and via its zinc finger domain, CDCA7 can interact with nucleosomal DNA (Unoki et al. 2019). Therefore, reduced levels of Cdca7 as observed upon Zbtb24 disruption, could lead to a hindrance for DNA methyltransferases to gain access to their genomic targets resulting in DNA hypomethylation.

The DNA methylation gain that we observed in the absence of Zbtb24 was intriguing, and since Zbtb24 was shown to preferably bind to promoters of actively transcribed genes (Thompson et al. 2018; Ren et al. 2019), we considered that its binding could influence DNA methylation levels *in cis*. By intersecting WGBS and ChIP-seq datasets we show that at a selected number of loci, Zbtb24 occupancy can indeed protect from aberrant DNA methylation gain. This is consistent with reports that local binding of transcription factors can contribute to maintain hypomethylated states of CGI (Krebs et al. 2014; Heberle and Bardet 2019). However, our results also show that loss of Zbtb24 binding is not always a predictor of hypermethylation, and very likely multiple layers of regulation exist that protect from aberrant hypermethylation. Likely, locus specific effects on DNA methylation patterns can be influenced by the sum of proteins recruited to that specific site (Blattler and Farnham 2013). We found that remaining H3K4me3 levels presented an obstacle for spreading of DNA methylation throughout a promoter CGI. This is in agreement with previous reports that this mark is crucial for the establishment and maintenance of unmethylated CGI at promoters because it repels DNA methylation (Weber et al. 2007). At CGI borders, where H3K4me3 levels are low, we observed DNA methylation gain upon loss of Zbtb24. It has been proposed that methylation changes at CGI shores are a function of TF binding (Krebs et al. 2014), and our observations are in agreement with such a scenario.

Together, our results suggest that there are at least two mechanisms by which Zbtb24 can influence DNA methylation and gene expression levels in mESCs. First, through an yet unknown pathway Zbtb24 helps to maintain DNA methylation at intergenic regions and gene clusters that despite DNA hypomethylation, do not get de-repressed in Zbtb24 mutant mESCs. Second, through its transcription factor function and direct binding to the promoters of actively transcribed genes, Zbtb24 regulates gene expression of its targets and helps to maintain the unmethylated state of CGIs. Here, Zbtb24 loss can result in transcriptional repression and aberrant DNA methylation gain. Relevant to this, mutations in ZBTB24 underlie ICF syndrome, a disease characterized by alterations in the DNA methylation landscape (Vukic and Daxinger 2019). Intriguingly, ZBTB24 has thousands of binding sites in somatic HCT116 cells and they are enriched for CpG-rich promoters of actively transcribed genes (Thompson et al. 2018). It will be interesting to determine how ZBTB24 ablation affects promoter DNA methylation and gene expression levels in differentiated tissues relevant to ICF syndrome pathology.

In summary, our study provides further insight on Zbtb24 function in DNA methylation regulation, and informs on how transcription factor binding contributes to the locus-specific regulation of gene expression patterns

## METHODS

### Mouse embryonic stem cell (mESC) culture, CRISPR/Cas9 genome editing and mESC transfection

Derivation of the *Zbtb24*^*+/+*^ and *Zbtb24*^*ΔBTB/ΔBTB*^ (C57/BL6) mouse embryonic stem cell (mESC) lines is described in (Wu et al. 2016). The *Zbtb24*^*-/-*^ E14 mESC knock out line was generated using CRISPR/Cas9 genome editing, and E14 cells were cultured on MEF feeders in serum conditions (Knockout DMEM (10829-018; Gibco), 10% FBS (DE14-801F; BioWhittaker), NEAA (11140; Gibco), L-Glutamine (25030-123; Gibco), Sodium Pyruvate (11360; Gibco), 2-Mercaptoethanol (31350; Gibco) and Leukemia Inhibitory Factor (ESG1107; Millipore)). *Zbtb24*^*+/+*^ and *Zbtb24*^*ΔBTB/ΔBTB*^ mESCs were cultured in serum conditions on 0.1% gelatin (G-1890; Sigma). For both 3xTy1-Zbtb24 overexpression and CRISPR/Cas9 transfections, Lipofectamine 3000 (L3000008; Thermo) was used following the online protocol. To generate *Zbtb24* knockout mESCs, vector pSpCas9(BB)-2A-Puro (PX459) V2.0 (a gift from Feng Zhang, Addgene plasmid #62988; http://n2t.net/addgene:62988; RRID: Addgene_62988) was used. The vector was cut with BpiI (ER1011; Thermo) and ligated with sgRNA annealed oligos for *Zbtb24*. 24 hours after transfection, cells were selected on puromycin (1µg/mL, P8833-10MG; Sigma) for 48 hours. Single colonies were picked and expanded for isolation of genomic DNA and characterized by Sanger sequencing. Potential knockout clones were then expanded for isolation of RNA and protein for further characterization by RT-qPCR and Western Blot. GuideRNA sequences and primers for genotyping are provided in Table S6.

### Genomic DNA isolation

Genomic DNA was isolated using the salt-extraction method. Briefly, cells were lysed in cell lysis buffer (50 mM Tris-HCl (pH 8), 4 mM EDTA (pH 8), 2% SDS) plus Proteinase K (390973P; VWR), and incubated at 55°C overnight. The next day, cell lysate was treated with RNaseA (EN0531; Thermo) at 37°C for 1 h. Saturated NaCl buffer was added followed by addition of isopropanol to precipitate genomic DNA and washing with 70% EtOH. Genomic DNA was dissolved in water and concentration was measured using Nanodrop.

### Molecular cloning

To generate the 3xTy1-Zbtb24 construct, full-length mouse *Zbtb24* (NM_153398.3) was PCR amplified from mESC cDNA library. All primers used for cloning can be found in Table S6.

### Western blot

Cells were lysed in Cell Lysis buffer (20 mM triethanolamine (T1377; Sigma), 0.14 M NaCl, 0.1% Sodium deoxycholate (D6750; Sigma), 0.1% SDS, 0.1% Triton X-100) with Protease Inhibitor Cocktail (27368400; Roche) and Phosphatase Inhibitor Cocktail (04906837001; Roche) on ice. BCA kit (23225; Thermo) was used to measure protein concentration. Equal amounts of total cell extracts were loaded on a NuPAGE gel (4–12%, NP0321; Thermo), and transferred to a Nitrocellulose Blotting Membrane (10600016; Life Sciences). The following primary antibodies were used: Zbtb24 (PM085; MBL Life Science, 1:1000), Cdca7 (15249-1-AP; Proteintech, 1:500), Dnmt3a (ab13888; Abcam, 1:1000), Dnmt3b (Ab16049; Abcam, 1:1000), Hells (11955-1-AP; Proteintech, 1:1000), GST (27457701; GE-Healthcare, 1:2000) and Tubulin (T6199; Sigma, 1:5000). Donkey anti-Rabbit 800CW (926-32213; Li-Cor, 1:5000), Goat anti-Rabbit 800CW (926-32211; Westburg, 1:5000), Donkey anti-mouse 680RD (926-68072; Li-Cor, 1:5000), Donkey anti-guinea pig 800CW (925-32411; LI-COR, 1:5000) were used as secondary antibodies. Membranes were analyzed on Odyssey (Westburg).

### RNA-seq *Zbtb24*^*+/+*^ and *Zbtb24*^*ΔBTB/ΔBTB*^ mESCs

Total RNA was isolated as described above and sample preparation was performed using NEBNext Ultra Directional RNA Library Prep Kit for Illumina (E7420S/L; NEB) according to the protocol. Libraries were sequenced with 125bp pair-end (PE) reads on a HiSeq 2500 at GenomeScan..

### Chromatin immunoprecipitation

Chromatin immunoprecipitation was described previously (Wu et al. 2016). Briefly, cells were cross linked with 1% formaldehyde (344198; Calbiochem) for 10 min at room temperature and glycine (125 mM) was used to quench cross-linking for 5 min. For mESCs, cells were washed twice with cold PBS and lysed with ChIP Buffer A (50 mM HEPES-KOH (pH 7.3), 140 mM NaCl, 1 mM EDTA (pH 8.0), 10% glycerol, 0.5% NP-40, 0.25% Triton X-100 and Protease Inhibitor Cocktail (05056489001; Roche)) for 10 min on ice. Samples were then spin down at 1,400 g for 5 min at 4°C, and the pellets were resuspended in ChIP Buffer B (1% SDS, 50 mM Tris-HCl (pH 8.0), 10 mM EDTA and Protease Inhibitor Cocktail) for 10 min on ice. For U2OS, cells were washed twice with cold PBS and lysed with NP Buffer (150 mM NaCl, 50 mM Tris– HCl (pH 7.5), 5 mM EDTA, 0.5% NP-40, 1% Triton X-100, Protease Inhibitor Cocktail). Lysed samples were sheared by sonication (Diagenode Biorupter Pico). Sheared chromatin was centrifuged at 12,000 g for 10 min at 4°C to discard the pellets. Before use, the supernatant of mESCs was diluted 10 times with NP buffer to make the final SDS concentration lower than 0.1%. For Ty1 ChIP, protein A and G Beads (10002D, 10003D; Life Technologies) were first blocked with 5mg/ml BSA (A7906; Sigma) and then incubated with antibodies at 4°C for at least 4 h. About 5 μg Ty1 antibody (C15200054; Diagenode), mouse IgG (12-371; Millipore) coupled with beads were incubated with sheared chromatin at 4°C overnight. For histone ChIP, chromatin was first incubated with 5 μg antibodies H3K4me3 (17-614; Millipore) or rabbit IgG (PP64; Millipore) at 4°C overnight. Protein A Sepharose beads (GE17528001; Sigma) blocked with 1mg/mL BSA (10848; Affymetrix) were added to pull down antibody-chromatin complex. After immunoprecipitation, beads were washed with low-salt washing buffer (0.1% SDS, 1% Triton X-100, 2 mM EDTA, 20 mM Tris-HCl (pH 8.1), 150 mM NaCl), high-salt washing buffer (0.1% SDS, 1% Triton X-100, 2 mM EDTA, 20 mM Tris-HCl (pH 8.1), 500 mM NaCl), LiCl washing buffer (0.25 M LiCl, 1% NP40, 1% deoxycholate, 1 mM EDTA, 10 mM Tris-HCl (pH 8.1)) and TE buffer (10 mM Tris-HCl (pH 8.0), 1 mM EDTA). Input DNA samples were extracted with phenol-chloroform-isoamylalcohol (15593049; Fisher Scientific). The immunoprecipitated DNA was purified using phenol-chloroform-isoamylalcohol, and concentration of pulled down DNA was measured using Qbit. For 3xTy1-Zbtb24 ChIP-seq in E14 mESCs, one input and one IP sample were sequenced by BGI on a BGISEQ-500 with 50 bp single-end (SE) reads.

### Quantitative ChIP-seq for H3K4me3 in mESCs

Cells were cross linked with 1% formaldehyde (M134-200ML; VWR) for 8 min at room temperature and glycine (125 mM; G8790-1KG; Sigma) was used to quench cross-linking for 5 min. Cells were washed twice with cold PBS and lysed in NP Buffer (150 mM NaCl, 50 mM Tris–HCl (pH 7.5), 5 mM EDTA, 0.5% NP-40, 1% Triton X-100, Protease Inhibitor Cocktail (05056489001; Roche)). Nuclei were sheared by sonication (Diagenode Biorupter Pico). For each H3K4me3 ChIP-seq experiment, 25 µg of sample chromatin was mixed with 50 ng spike-in *Drosophila* chromatin (53083; Active Motif). Mixture of experimental chromatin and spike-in chromatin was then incubated with a mix containing 4 µg of H3K4me3 antibody (CS200580; Millipore) and 2 µg of spike-in antibody (104597; Active Motif) at 4°C overnight. The next day, Protein A Sepharose beads (175280-01; GE Health Care) were first blocked with 1mg/ml BSA (10484; Affymetrix) and then added to each chromatin-antibody mix and incubated at 4°C for at least 3 h. After immunoprecipitation, beads were washed with low-salt washing buffer (0.1% SDS, 1% Triton X-100, 2 mM EDTA, 20 mM Tris–HCl (pH 8.1), 150 mM NaCl), high-salt washing buffer (0.1% SDS, 1% Triton X-100, 2 mM EDTA, 20 mM Tris–HCl (pH 8.1), 500 mM NaCl), LiCl washing buffer (0.25 M LiCl, 1% NP40, 1% deoxycholate, 1 mM EDTA, 10 mM Tris–HCl (pH 8.1)) and TE buffer (10 mM Tris–HCl (pH 8.0), 1 mM EDTA). DNA was extracted using phenol-chloroform-isoamylol (15593-049; Life Technologies). Samples were sequenced at GenomeScan on a NovaSeq6000 with 150 bp paired-end (PE) reads.

## QUANTIFICATION AND STATISTICAL ANALYSIS

### Statistical analyses

A Student’s test and Standard error of mean (SEM) were used for all the statistical analysis from at least two biological replicates or two independent experiments. Values of *p* <0.05 were considered to be significant. A Fisher exact test implemented in BEDTools v2.28 was used to for annotation enrichment. To test global methylation differences, a paired samples Wilcoxon rank sum test was selected. All tests were two-sided unless stated otherwise.

### RNA-seq analysis

Quality assessment of the raw sequencing reads was done using FastQC v0.11.2 (http://www.bioinformatics.babraham.ac.uk/projects/fastqc). Adapters were removed by TrimGalore v0.4.5 (https://www.bioinformatics.babraham.ac.uk/projects/trim_galore/) using default parameters for paired-end Illumina reads, after which, quality filtering was performed by the same software. Reads smaller than 20bps and those with an error rate (TrimGalore option “-e”) higher than 0.1 were discarded, after which a final quality assessment of the filtered reads was done with FastQC to identify possible biases left after filtering. The remaining reads were mapped to the mouse reference genome (build mm10) using the STAR aligner v2.5.1(Dobin et al. 2013) using default parameters with the following exceptions: “– outputMultimapperOrder random” and “–twopassMode basic”. Before mRNA quantification, duplicated reads were marked with Picard tools v2.17 (http://broadinstitute.github.io/picard/).

Quantification was done by HTSeq-count v0.91 (Anders et al. 2015), using the GENCODE MV16 annotation with the option “–stranded no”. Statistical analysis was done using DESeq2 v1.2.0 (Love et al. 2014) (R package). Figures 5A-C were created with custom R scripts available via (https://git.lumc.nl/dsanleongranado/zbtb24_figures). The final list of differential expressed genes contains genes for which the adjusted *p*-value (Benjamini-Hochberg correction) <0.05 and |log_2_(FC)| >1. The quantification of exon expression was generated with SGSeq v1.17.0 (Goldstein et al. 2016) (R/Bioconductor package) and the differential exon usage analysis was done with DEXSeq (Anders et al. 2012).

### RRBS analysis

The preprocessing and deduplication of the reads was done following the workflow described by NuGen Technologies (https://github.com/nugentechnologies/NuMetRRBS). For the methylation level calculation, only CpGs with a coverage higher than 5 were taken into account. These methylation levels were used to identify differentially methylated regions (DMRs) between *Zbtb24*^*+/+*^ and *Zbtb24*^*ΔBTB/ΔBTB*^ samples. DMR calling was done by DSS, version 2.28 (Feng et al. 2014) (R/Bioconductor package). For downstream analysis, only the regions with a *p*-value <.05, a minimum number of CpGs greater or equal than 7 and a minimum difference of methylation of >15% were taken into account. The annotation to genes was performed using ChIPseeker v1.18.0 (Yu et al. 2015) (R/Bioconductor package), the promotor region was defined as TSS ±2kb.

### WGBS analysis

Raw sequences were filtered by quality with TrimGalore using default parameters. Reads with length smaller than 20 bps and error rate higher than 0.1 were discarded. After which, the sequences were aligned to mm10 using the Bismark aligner v0.18 (Krueger and Andrews 2011), to increase sensitivity we used parameter “-N 1”. The duplicates in the alignment were removed with Deduplicate Bismark(part of the Bismark package). Methylation calling was performed by Bismark Methylation Extractor (part of the Bismark package), using default parameters with the following exceptions: “–paired-end”, “–ignore_r2 2” and “–bedgraph”. For DMR calling we used the algorithm described in (Wu et al. 2015), to select the single-replicate algorithm in DSS v2.31.0 (R/Bioconductor package) (Feng et al. 2014), the parameter “smoothing=TRUE” was included. For downstream analysis, the list of DMRs was filtered for *p*-values less than 0.05, number of CpGs greater or equal than 7 and a minimum difference of methylation greater than 15. The region annotation was done by HOMER v4.10 (Heinz et al. 2010) and the sample correlations were calculated with Methylkit v1.0.9 (Akalin et al. 2012). DMR enrichment analysis was done using the annotations provided by annotatePeak from ChIPseeker R package. The statistical analysis was done with BEDTools fisher with default parameters and mm10 chromosome sizes.

### Genome-wide DNA methylation analysis for repetitive elements

Methylation calls provided by Bismark Methylation extractor for the reads mapped uniquely to the mouse genome were intersected with UCSC RepeatMasker track (mm10). Overlapping reads and their respective methylation percentage were annotated, sorted and grouped by the repetitive element subfamilies, repetitive element families and repetitive classes using bedtools v2.28 (Quinlan and Hall 2010). DNA methylation levels of the interspersed repeats for the repeats-superfamilies, satellites, LINEs, ERVs and SINEs were plotted using the R packages pheatmap v1.0.12 and ggplot2 v3.1.1.

### Histone modification ChIP-seq analyses

The H3K4me3, H3K9me3, H3K36me3 and H3K27me3 mESC ChIP-seq datasets were downloaded from GEO (GSE23943) (Marks et al. 2012). The reads were preprocessed with TrimGalore to remove low quality reads, the resulting reads were aligned to the mouse genome (mm10) using bowtie2 v2.3.4.2 (Langmead and Salzberg 2012) with default parameters. To generate the tracks and the heatmap and profile plots, deepTools version 3.1.3 (Ramirez et al. 2016) was used. For peak calling of H3K4me3, MACS2 v2.1.0 (Zhang et al. 2008) was used with the following settings: “-g mm”, “–B” and “-qvalue 0.05”. For peak calling of H3K9me3, H3K36me3 and H3K27me3 modifications “--broad” was included. For the plot profiles, a bin size of 50 bps and a smoothing window of 100 bps was chosen. To remove outliers (regions with very high read counts), the parameter “maxThreshold” was set to 20. The data visualization was done with in-house R scripts (https://git.lumc.nl/dsanleongranado/zbtb24_figures) using Gviz v1.28.0 (Hahne and Ivanek 2016) (R / Bioconductor package), the plotted tracks were smoothed using a 50 bps sliding window. For downstream analysis, the CpG island dataset for the mouse genome (mm10) was downloaded from UCSC (https://genome.ucsc.edu/, accessed on 15/11/2018).

### H3K4me3 spike-in ChIP-seq in *Zbtb24*^*+/+*^ and *Zbtb24*^*ΔBTB/ΔBTB*^ mESCs

The preprocessing of the samples follow the same steps described below (see Section Histone ChIP-seq analysis). To reduce the effects of technical variation and sample processing bias, the datasets were Spike-in normalized (Egan et al. 2016). To obtain the normalizing factors, the reads were mapped to the mouse genome (build mm10) and the *Drosophila melanogaster* genome (build dm6) with the Bowtie2 aligner using the “–very-sensitive” parameter. Duplications were removed with the Picard toolkit and the multiple mapped reads and low quality alignments were filtered out using SAMtools v1.9 (Li et al. 2009) using a mapping quality threshold (MAPQ) > 30. The spike-in factors were calculated using the remaining reads as follows:

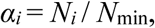

with *N*_*i*_ the number of unique alignments for sample *i* and *N*_min_ the minimum number of unique alignments in all samples. Spike-ins were used to down sample the datasets with the down sample function of Picard tools, which randomly remove read pairs with a probability equal to the reciprocal of the spike-in factor. The normalized samples were used to identify differential binding events between *Zbtb24*^*+/+*^ and *Zbtb24*^*ΔBTB/ΔBTB*^ mESCs. This was performed with the sliding window approach implemented in PePr v1.1.10 (Zhang et al. 2014) with default parameters with the exception of “–threshold=0.05” and “–normalization no”. For visualization and downstream analysis, the sample correlations and bigWig files creation was done with deepTools with a bin size of 100 bps.

### Analysis of mESC Ty1-Zbtb24 ChIP-seq

For ChIP-seq analysis, reads were mapped to the mouse reference genome (mm10) using BWA-mem v0.7.16 (Li and Durbin 2010) with default parameters. After alignment, reads with low mapping quality (MAPQ < 30) were filtered out using SAMtools. For peak calling and bigWig track generation, MACS2 v2.1.0 (Zhang et al. 2008) was used with the following settings: “-g hs”, “–B” and “-qvalue 0.05” for U2OS and “-g mm”, “-B” and “-qvalue 0.05” for mESC. Input datasets containing the whole genomic DNA fragments were used as controls. The Zbtb24 motif was identified using MEME-ChIP v5.0.3 (Machanick and Bailey 2011) with default settings. Reads overlapping with the intervals of Zbtb24 binding sites in mESCs were extracted with bedtools. The heatmap was generated using pheatmap v1.0.12 (R package; https://cran.r-project.org/web/packages/pheatmap/index.html).

## DATA ACCESS

The data used in this study have been deposited in the NCBI Gene Expression Omnibus (GEO; http://www.ncbi.nlm.nih.gov/geo/) under accession code GSE131260.

## ACKNOWLEDGEMENTS

We thank S. vd Maarel, B. Heijmans, D. Sikrova and V. Della Chiara for helpful discussions and suggestions on the manuscript. We thank S. Kloet, E. de Meijer and Y. Ariyurek for help with NGS sample preparation and I. Karemaker for advice on quantitative ChIP-seq. This work was financed through grants from the Leiden University Medical Centre (LUMC Fellowship) and the Netherlands Organization for Scientific Research (NWO-ZonMW-Vidi 91718350) to L.D.

## AUTHOR CONTRIBUTIONS

H.W. contributed to study design, performed experiments, analyzed and interpreted data and contributed to bioinformatics analyses. D.S.L.G. performed bioinformatics analyses and statistical analyses. M.V. contributed to study design, performed experiments and analyzed and interpreted data. K.K.D.V. and C.B. performed experiments and analyzed data. J.C. performed bioinformatics analyses and interpreted data. J.F.J.L. contributed to data interpretation, bioinformatics and statistical analyses. L.D. conceived and supervised the study, interpreted data and wrote the manuscript with input from all authors.

---

**Figure S1.**
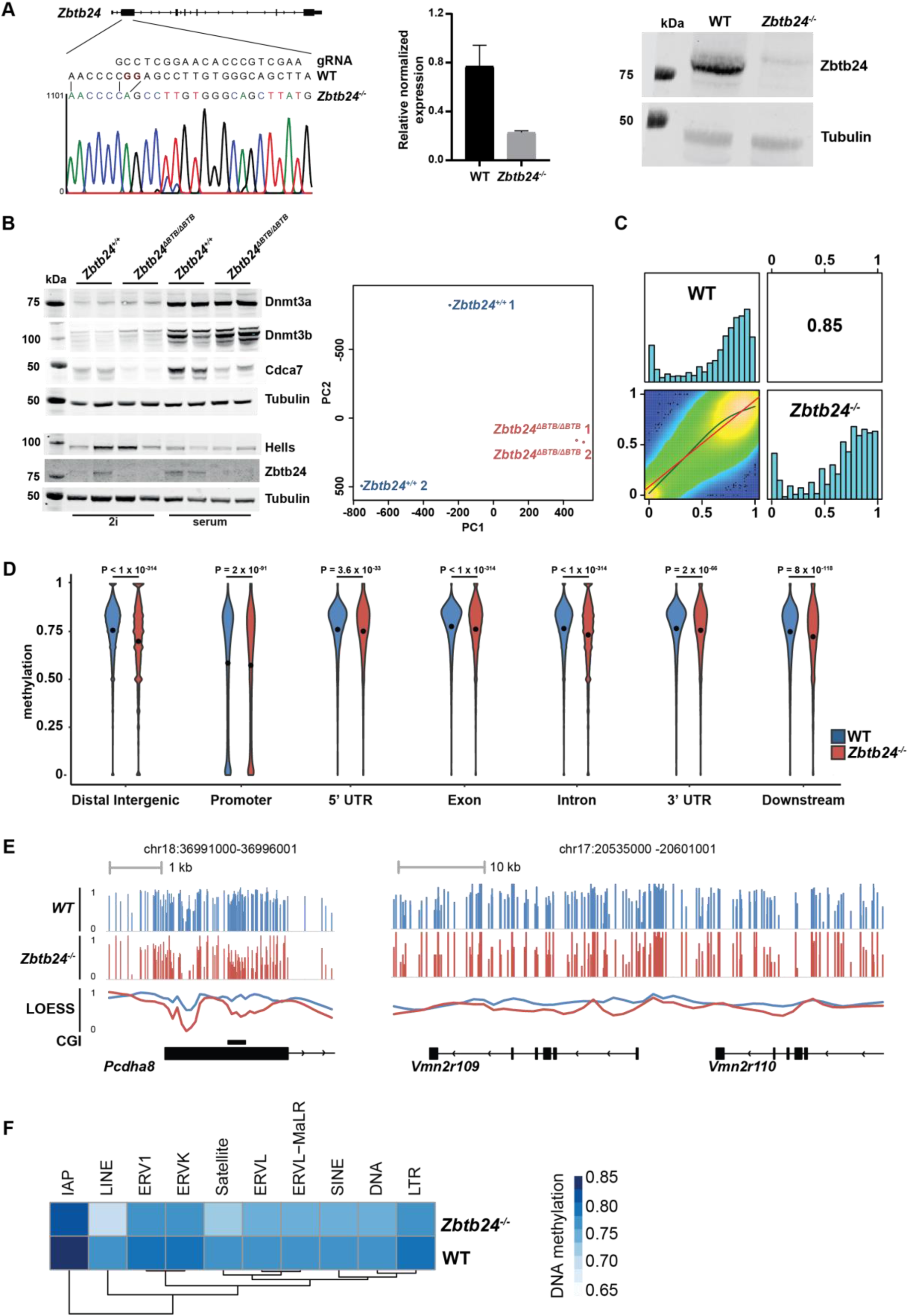
Generation of the *Zbtb24*^*-/-*^ CRISPR/Cas9 knock out mESC line and DNA methylation changes in *Zbtb24*^*-/-*^ and *Zbtb24*^*ΔBTB/ΔBTB*^ mutant mESCs. (**A**) Generation of *Zbtb24*^*-/-*^ CRISPR/Cas9 knock out mESCs. (left) Sanger sequencing traces showing genomic DNA of the *Zbtb24*^*-/-*^ mESC line carrying a homozygous 2 nt deletion in exon 2 of the *Zbtb24* gene, resulting in a premature stop codon. (middle) RT-qPCR showing reduced *Zbtb24* mRNA levels. (right) Western blot showing severely reduced Zbtb24 protein in the *Zbtb24*^*-/-*^ mESC line. Tubulin was used as a loading control. **(B)** (left) *Zbtb24*^*+/+*^ and *Zbtb24*^*ΔBTB/ΔBTB*^ mESCs were switched from 2i to serum conditions for RRBS analysis. Western blot depicting protein levels of Dnmt3a/b, Cdca7, Hells and Zbtb24 in *Zbtb24*^*+/+*^ and *Zbtb24*^*ΔBTB/ΔBTB*^ mESCs cultured either in 2i or serum conditions. Tubulin was used as a loading control. (right) PCA plot of principal components 1 and 2 of RRBS analysis showing clustering of *Zbtb24*^*+/+*^ and *Zbtb24*^*ΔBTB/ΔBTB*^ mESC samples. (**C**) Scatter plot and correlation of CpG methylation between WGBS samples. Heat plots show pairwise comparisons of methylation levels for the two samples. Numbers in the upper right corner denote Pearson correlation coefficients. Histograms on the diagonal show methylation level frequency per cytosine for each sample. (**D**) Violin plots showing methylation levels (3kb genomic tiles) of genomic regions in WT (blue) and *Zbtb24*^*-/-*^ (red) WGBS samples. The data is aggregated by promoter (TSS±2kb), 5’UTR, exons, introns, 3’UTR, downstream regions (TES+2kb). Remaining CpGs are included in the distal intergenic class. The black dots represent the means. (**E**) UCSC screenshot depicting methylation levels (WGBS) of representative hypomethylated loci in *Zbtb24*^*-/-*^ mutant mESCs. The line plot represents the locally estimated scatter plot smoothing (LOESS) of WT (blue) and *Zbtb24*^*-/-*^ (red) methylation levels. **(F)** Unsupervised hierarchical clustering and heatmap of methylation average of repetitive elements across WT and *Zbtb24*^*-/-*^ mutant mESCs. Dark blue represents higher and light blue lower methylation levels.

**Figure S2.**
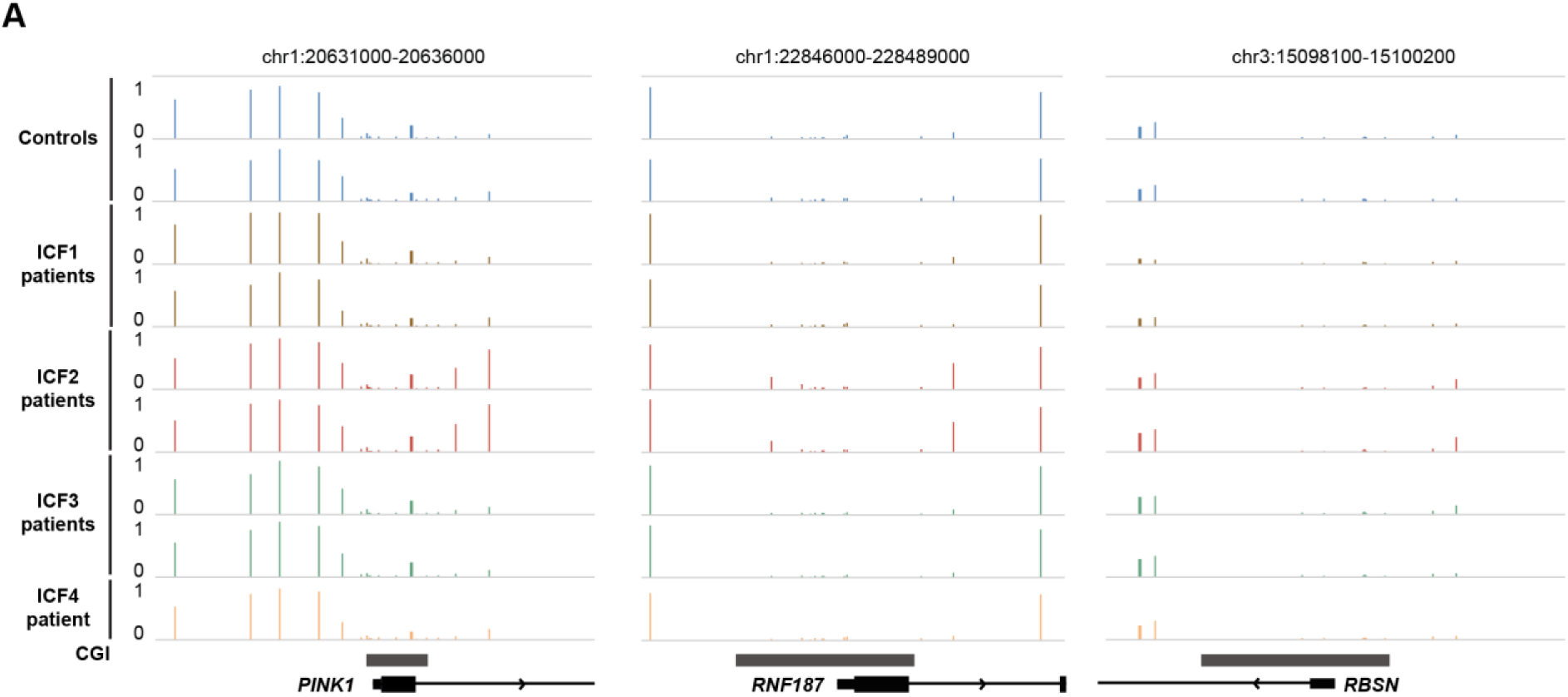
Representative UCSC genome browser screenshots. **(A)** UCSC genome browser screenshots showing DNA methylation levels (Velasco et al., HMG 2018; Illumina 450K array data) of genomic loci corresponding to Figure 1I, in control samples (blue), ICF1 patients (orange), ICF2 patients (red), ICF3 patients (green) and ICF4 patients (yellow). Grey rectangles represent CpG islands (CGI).

**Figure S3.**
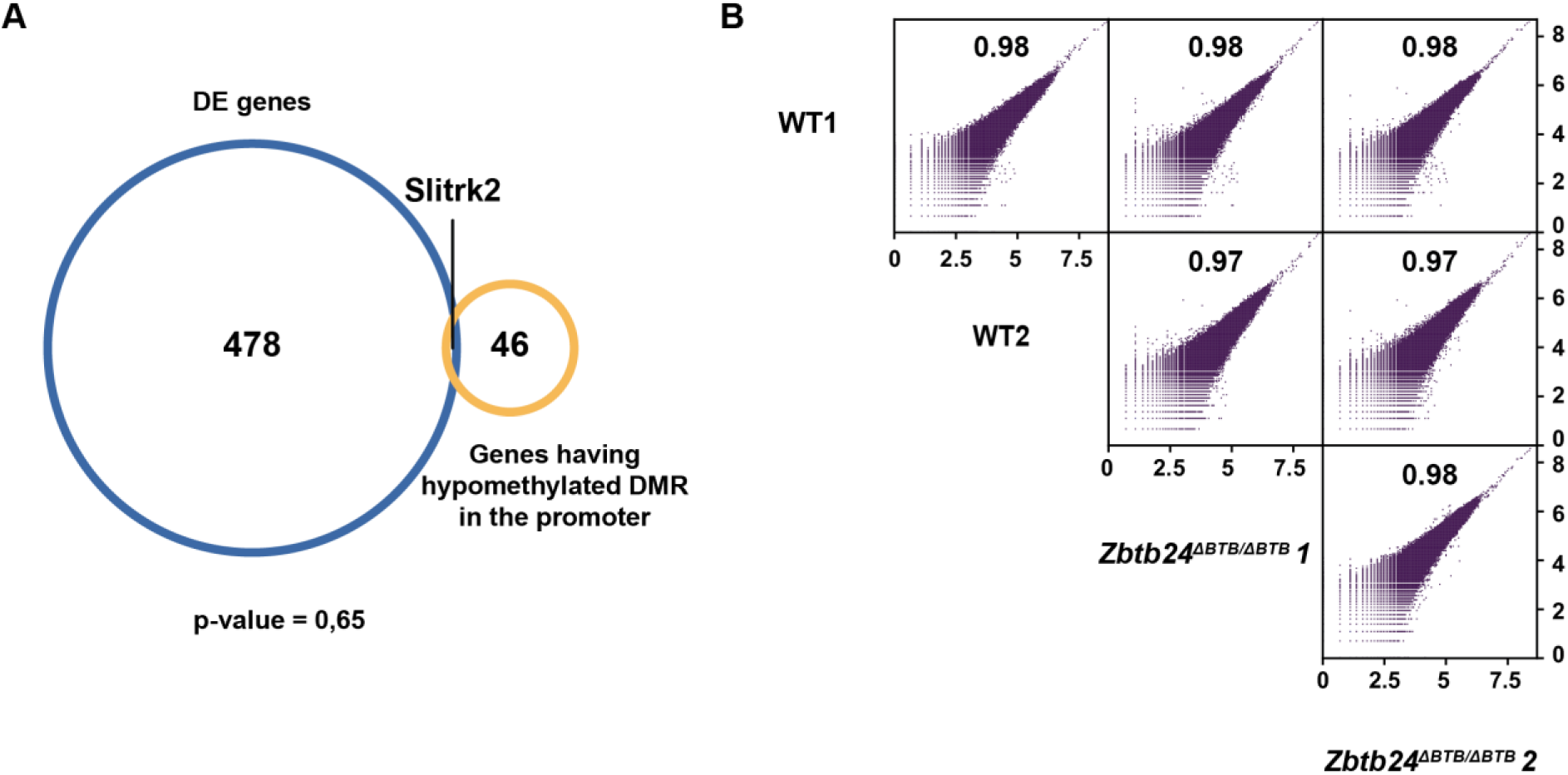
WGBS and RNA-seq overlap, and H3K4me3 ChIP-seq data. **(A)** Venn diagram showing the intersection between genes +2 kb upstream sequence that overlap with a hypomethylated DMR (*Zbtb24*^*-/-*^ versus WT) and differentially expressed genes (*Zbtb24*^*ΔBTB/ΔBTB*^ versus *Zbtb24*^*+/+*^). The *p*-value (hypergeometric test) indicates the significance of the overlap. Background was calculated using “expressed genes only (base Mean>5)”. **(B)** Scatter plot depicting the correlation of the H3K4me3 signal at 100 bp windows between two *Zbtb24*^*+/+*^ and two *Zbtb24*^*ΔBTB/ΔBTB*^ samples. The axis are log scaled and the numbers inside the plots represent the Pearson correlation.

